# Generation of a Sleeping Beauty transposon-based cellular system for rapid and sensitive identification of SARS-CoV-2 host dependency and restriction factors

**DOI:** 10.1101/2021.04.27.441606

**Authors:** Marek Widera, Alexander Wilhelm, Tuna Toptan, Johanna M. Raffel, Eric Kowarz, Fabian Roesmann, Anna Lena Siemund, Vanessa Luciano, Marius Külp, Jennifer Reis, Silvia Bracharz, Christiane Pallas, Sandra Ciesek, Rolf Marschalek

**Affiliations:** Institute for Medical Virology, University Hospital Frankfurt am Main, Goethe University, 60596 Frankfurt am Main, Germany; Institute of Pharmaceutical Biology, Goethe-University, 60438 Frankfurt/Main; German Center for Infection Research, DZIF, 60596 Braunschweig, Germany; Fraunhofer Institute for Molecular Biology and Applied Ecology (IME), Branch Translational Medicine and Pharmacology, 60596 Frankfurt am Main, Germany

**Keywords:** SARS-CoV-2, COVID-19, corona virus, reporter cell line, high throughput, drug screening, CPE

## Abstract

The severe acute respiratory syndrome coronavirus-2 (SARS-CoV-2) is the causative agent of the acute respiratory disease COVID-19, which has become a global concern due to its rapid spread. The common methods to monitor and quantitate SARS-CoV-2 infectivity in cell culture are so far time-consuming and labor-intensive. Using the Sleeping Beauty transposase system, we generated a robust and versatile reporter cell system that allows SARS-CoV-2 infection experiments compatible for high-throughput and live cell imaging. The reporter cell is based on lung derived A549 cells, which show a profound interferon response and convenient cell culture characteristics. ACE2 and TMPRSS2 were introduced for constitutive expression in A549 cells. Subclones with varying levels of ACE2/TMPRSS2 were screened for optimal SARS-CoV2 susceptibility. Furthermore, extensive evaluation demonstrated that SARS-CoV-2 infected reporter cells were distinguishable from mock-infected cells and already showed approximately 12 h post infection a clear signal to noise ratio in terms of cell roughness, fluorescence and a profound visible cytopathic effect. Moreover, due to the high transfection efficiency and proliferation capacity, Sleeping Beauty transposase-based overexpression cell lines with a second inducible fluorescence reporter cassette (eGFP) can be generated in a very short time, enabling the investigation of host and restriction factors in a doxycycline-inducible manner. Thus, the novel reporter cell line allows rapid and sensitive detection of SARS-CoV-2 infection and the screening for host factors essential for viral replication.

**Highlights:**

- Sleeping Beauty transposon-based cellular system was used to generate a highly susceptible cell line for monitoring SARS-CoV-2 infection
- The versatile reporter cell line A549-AT is suitable for rapid and sensitive high-throughput assays
- Additional gene specific expression cassettes allow the identification of SARS-CoV-2 host dependency and restriction factors

## Introduction

The coronavirus disease 2019 (COVID-19) is caused by an infection with the severe acute respiratory syndrome coronavirus 2 (SARS-CoV-2). The origin of SARS-CoV-2 outbreak was initially described in Wuhan, China (Zhu et al., 2020) and rapidly established a worldwide pandemic. Globally, over 135 million cases and 2.9 million deaths in total since the start of the pandemic have been reported by the WHO as published on April, 11^st^ 2021 (WHO, 2021).

SARS-CoV-2 is a spherical beta coronavirus with a size of 120 nm in diameter, which has a lipid envelope. Viral cell entry depends on the host receptors carboxypeptidase angiotensin I converting enzyme 2 (ACE2) and the cellular transmembrane protease, serine 2 (TMPRSS2) for the priming of the viral glycoprotein (Hoffmann et al., 2020b; Rahman et al., 2020). ACE2 was already identified a functional receptor for SARS-CoV (Li et al., 2003) and physiologically plays a key role in the renin-angiotensin aldosterone system by its ability to degrade angiotensin II to the metabolites angiotensin 1–7 and 1-9 (Keidar et al., 2007; McKinney et al., 2014). The spike glycoproteins (S) of SARS-CoV and SARS-CoV-2 receptor-binding domain (RBD) bind to the ACE2 receptor (Lan et al., 2020). Albeit SARS-CoV-2 RBD was shown to have a higher ACE2 binding affinity compared to SARS-CoV RBD, the entire SARS-CoV-2 S binding affinity is not higher when compared to SARS-CoV, suggesting that SARS-CoV-2 RBD, albeit more potent, is less exposed than SARS-CoV RBD (Shang et al., 2020; Wang et al., 2020b).

Activation of the S protein and viral entry is further mediated via sequential proteolytic cleavage by TMPRSS2 enabling the glycoprotein mediated membrane fusion of the viral envelope with the host cell membrane (Belouzard et al., 2009; Heurich et al., 2014). Of note, cells infected with SARS-CoV, MERS-CoV, or SARS-CoV-2 were shown to express S protein on the cell surface and are able to induce syncytia and the formation of a morphological cytopathic effect (CPE) (Buchrieser et al., 2020; Bussani et al., 2020; Chan et al., 2013; Hoffmann et al., 2020a; Matsuyama et al., 2010; Qian et al., 2013). Furthermore, TMPRSS2 was shown to facilitate and accelerate syncytia formation by promoting the fusion process (Buchrieser et al., 2020).

After entry, the viral genomic RNA (vRNA), which serves as a template for initial polyprotein translation and generation of non-structural proteins 1-16 (NSP1-16), is subjected to complementary transcription catalyzed by the viral RNA dependent RNA-polymerase (RdRP). In addition, the full genomic repertoire is expanded by the usage of a process named discontinuous transcription in which subgenomic RNAs (sgRNAs) are formed (Kim et al., 2020) serving as templates for the translation of downstream ORFs 3a-9b, including the E, M, and N genes. SgRNAs are each composed of an RNA primer that can co-transcriptionally jump to sequences at transcription-regulating sequences (TRSs) and create a junction with downstream sequence elements that code for the other viral proteins. Quantitative reverse transcriptase polymerase chain reaction (RT-qPCR) is the commonly used method for sensitive and specific detection method for SARS-CoV-2. In particular, targeting the vRNA derived from cell culture supernatants is suitable for the quantification of genome copy equivalents (Toptan et al., 2020) while the detection of specific intracellular sgRNAs is used to quantify active viral replication (Kohmer et al., 2021; Shin et al., 2020; Wolfel et al., 2020). However, high costs, excessive hands on time, and reagent shortages emerging during the pandemic disqualify RT-qPCR for high-throughput testing.

*In vitro* cell culture models that can realistically mimic the viral replication cycle to decipher the pathology of COVID-19 are limited.

Primary human airway epithelial cells highly express both receptors ACE2 and TMPRSS2 and are permissive for SARS-CoV-2. They show cytopathic effects 96 h post infection (Hoffmann et al., 2020b; Takayama, 2020), but have limited lifespan and thus difficult to handle or expensively available from commercially sources. Currently, the common cell lines for SARS-CoV-2 research are Caco2, Calu-3, Vero E6, HEK293T, and Huh7. Vero E6 cells have been shown to be susceptible to SARS-CoV-2 (Ogando et al., 2020). By introducing additional gene copies of TMPRSS2 they have been rendered even superior to infection when compared to the parental cell line by 2-log (Matsuyama et al., 2020). However, a severe impairment to work with these cells and to draw conclusions about the interaction with the host is only possible to a very limited extent, since an essential component of the type I interferon signaling pathway is defective (Osada et al., 2014).

Caco2 are highly susceptible to SARS-CoV-2 and commonly used in our lab, showing a clear cytopathic effect (Bojkova et al., 2020b; Hoehl et al., 2020; Shin et al., 2020; Toptan et al., 2020). However, working with these human colon adenocarcinoma derived cells is very struggling, since CaCo2 have suboptimal growth properties in cell culture and inefficient transfection and selection properties and, thus, are not suitable for high throughput analysis.

In addition, two human lung epithelial cell lines, A549 (alveolar epithelial cell line) and Calu-3 (bronchial epithelial cell line) are available. Calu-3 cells are susceptible to SARS-CoV-2 infection (Bestle et al., 2020; Bojkova et al., 2020a) with a comparable efficiency when compared to Caco2 cells (Stanifer et al., 2020). In contrast to Calu-3, A549 cells have superior culturing properties compared to Caco2 cells and were shown to efficiently produce IFNγ (Torvinen et al., 2007) indicating an intact interferon chemokine signaling pathway. However, parental A549 cells are poorly infectable with SARS-CoV-2 (Hoffmann et al., 2020b).

The aim of this work was to establish a lung cell based system that would allow infection experiments in a cell line that is easy to maintain, suitable for large-scale propagation, and importantly, features an intact Type I / III and II signaling pathway. Furthermore, an inducible gene overexpression cassette, for the investigation of cellular host restriction or dependency factors should be implemented.

## Results

The major prerequisites for a novel reporter cell line for high throughput SARS-CoV-2 research are easy cultivation properties, reproducibility, relatively unlimited availability, but also pronounced interferon responses. As a suitable starting point, we have chosen the adenocarcinomic human alveolar basal epithelial cells A549 cells (Giard et al., 1973) and initially tested the susceptibility to interferons in comparison to in SARS-CoV-2 research commonly used Caco2 and Vero cells. Cells were stimulated with either type I or II interferons and the mRNA expression levels of known interferon stimulated genes (ISGs) were examined by RT-qPCR. As surrogate markers for ISGs, we have chosen the two genes Interferon-stimulated gene 15 (ISG15) and Interferon regulatory factor 1 (IRF1), which are predominantly stimulated by type I / III or II interferons, respectively, and are significantly involved in host cell defense against SARS-CoV-2 (Karki et al., 2021; Shin et al., 2020). Compared to Caco2 cells (24-fold), an approx. 10-fold higher expression of ISG15 was detected in A549 cells (275-fold) when treated with type I interferon (**Fehler! Verweisquelle konnte nicht gefunden werden.a**). In Vero cells no ISG15 expression was observed after IFN-I stimulation confirming the defect in IFN-I signaling (Osada et al., 2014). In all cell lines IFNγ treatment resulted in >2 log induction of IRF1 expression (345-fold in Caco2; 348-fold in A549 cells; 412-fold IRF1 increase in Vero cells). A certain crosstalk could also be observed, since IFNγ treatment of A549 and Caco2 cells resulted in a moderate but highly significant increase of ISG15 expression. Due to the defect in type I interferon signaling, Vero cells were excluded from further analysis.

Next, the time course of the ISG induction between A549 and Caco2 cells was investigated and two type III interferons (IFN-λ1 and -λ2), which have a distinct receptor to type I interferons but share common downstream signaling cascade, were also added. In both A549 and Caco2 cells, treatment with IFN-β (type I IFN) resulted in a strong increase with higher ISG15 expression levels, while both IFN-λ subtypes induced a similar increase and maximal level which were about one log level lower when compared with IFN-β treated cells. IRF1 expression could be induced most strongly by IFN-γ while a moderate expression could be measured after IFN-β stimulation. The maximum value was measured at 3 hours after stimulation. Stimulation with IFN-lambda resulted in marginal ISG expression only.

These data indicate that the A549 is suitable cell line to study type I, II, and III interferon dependent effects.

Since the choice of a cell line for high-throughput purposes depends largely on its properties in cell culture, we compared proliferation rate, trypsinization and transfection efficiency of A549 and Caco2 cells. For this purpose, cells were seeded with low density and the cell growth was monitored over a period of 16 days using an automated confluence measurement and cell counting system. Exponential proliferation in A549 was observed immediately after seeding reaching full confluency at day 8 (**Figure 2A**). Caco2 cells proliferated in a biphasic manner, with very slow growth observed for the first few days (until about day 4), reaching a 50% confluency at day 8, and approximately 75% at day 15. A further characteristic for efficient use is the duration of trypsinization until full detachment of the cells prior to splitting for experiments. A549 cells required a significantly shorter time than Caco2 cells, which were detached from a plate after around 3 min (**Figure 2B**). Finally, the transfection efficiency of the two cell lines was compared. A GFP-encoding plasmid DNA (pSBtet GP) was used for transient transfection of both cell lines. Using standard conditions, we found that A549 cells allowed a 70% higher transfection efficiency than Caco2 cells (**Figure 2C**). These data confirmed that A549 have cell culture properties that are highly suitable for high-throughput analyses.

**Figure 1.**
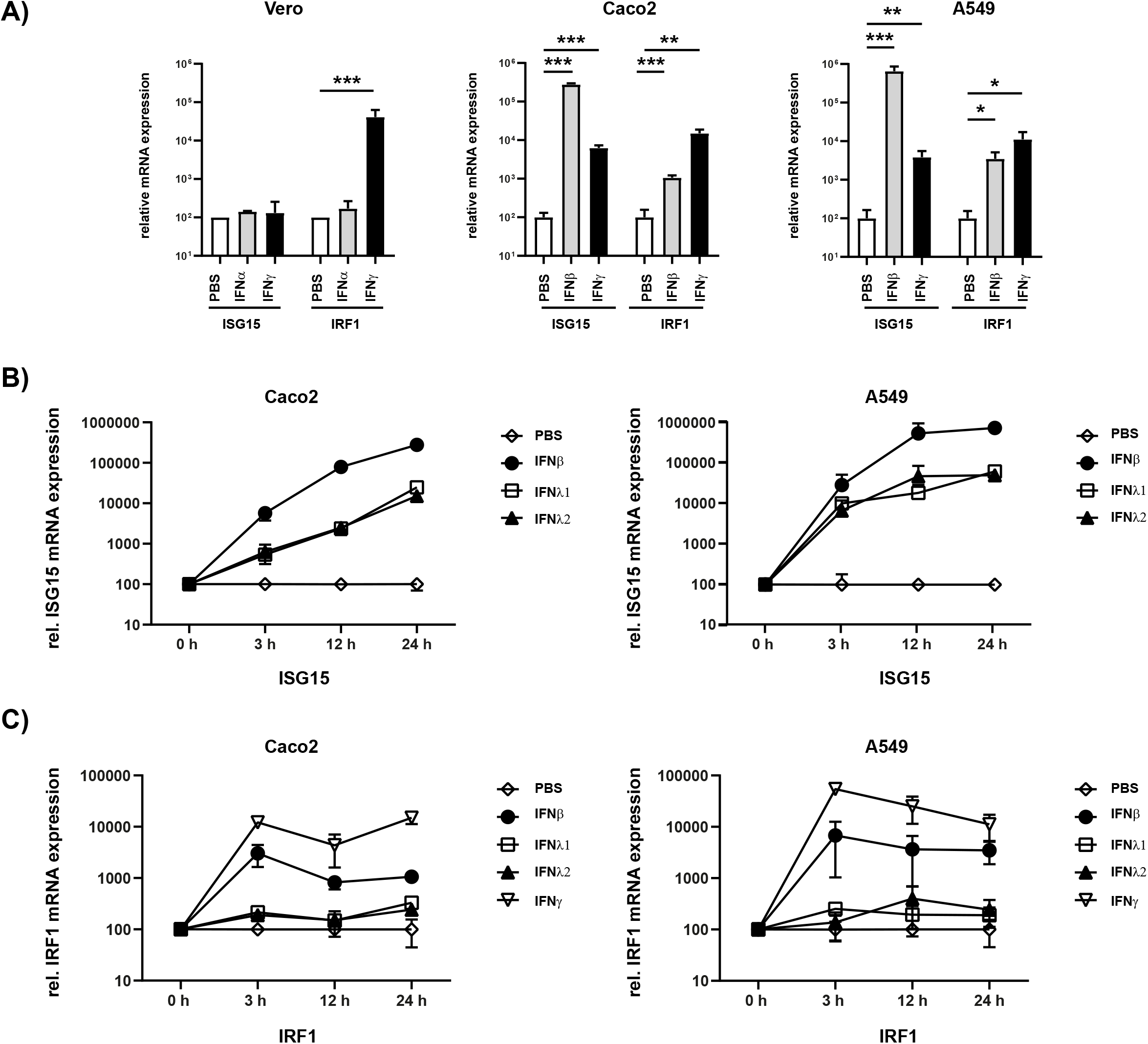
Susceptibility of cell lines to type I-III interferons. Vero, Caco2 and A549 cells were stimulated with the indicated type I/III (500 U / mL) and type II interferons (10 ng / mL). **A)** After 24 hours, total RNA was isolated and subjected to RT-qPCR analysis. ISG15 (under control of ISRE promoter) was used as a surrogate marker for IFN-I and III. Activation of the type II interferon pathway was monitored by IRF1 expression (under control of the GAS promoter). **B and C)** Time course of ISG15 and IRF1 gene expression in Caco2 and A549 cells. RNA was isolated at the indicated time point and subjected to RT-qPCR analysis. Expression was calculated using the Δct method and beta actin as housekeeping gene. Error bars indicate SD from three biological replicates. * (p < 0.05), ** (p ≤ 0.01) and *** (p ≤ 0.001).

**Figure 2.**
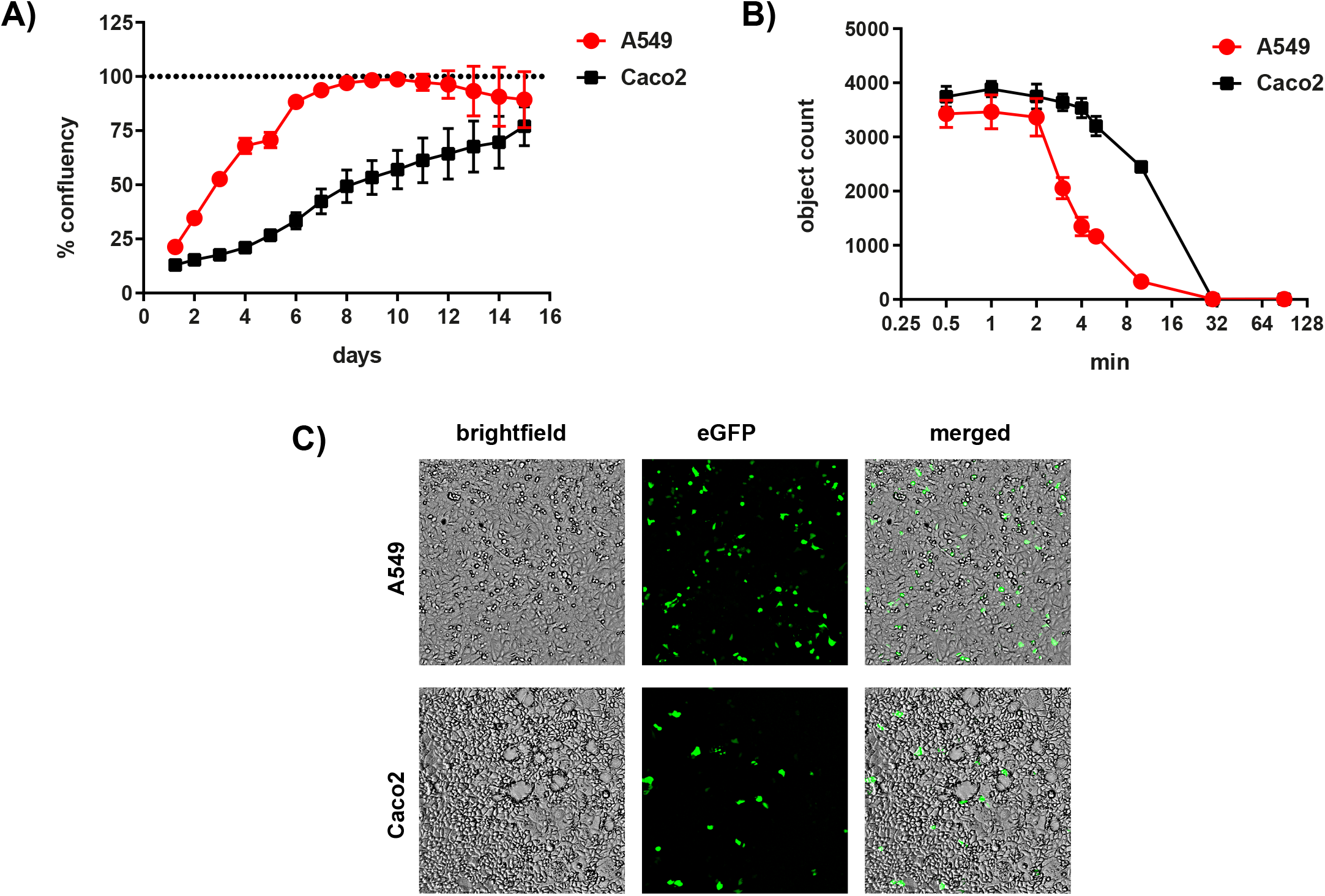
Cell culture characteristics of A549 and Caco2 cells. **A)** Growth curves of Caco2 and A549 cells were shown in 12-well plates with low density (2×10^4^ A549-AT / well; 1×10^5^ Caco2 / well). The relative confluence was monitored automatically using live cell imaging SparkCyto 400 (Tecan) over a period of 198 hours. After 48 hours, each well was washed with PBS to remove unattached cells. **B)** Trypsinization properties of A549 and Caco2 cells. Cells were seeded in 96-well plates, treated with trypsin/EDTA and incubated for the indicated time. For incubation times longer than 10 min cells were incubated at 37°C, otherwise at room temperature. **C)** To determine transfection efficiency, A549 and Caco2 cells were transiently transfected with an eGFP-encoding plasmid (pSBtet GP). After 48 h, eGFP fluorescence was detected.

Since parental A549 cells are less susceptible to SARS-CoV and SARS-CoV-2 infection and per se do not allow CPE formation, we aimed to overexpress both receptors ACE2 and TMPRSS2 (Hoffmann et al., 2020b) necessary for SARS-CoV-2 entry. We used the well-established Sleeping Beauty system (Kowarz et al., 2015), which enables transposase-mediated stable transfection of the target cells. For this purpose, we first generated a construct that allows constitutive expression of ACE2 together with an expression cassette for dTomato (**Fehler! Verweisquelle konnte nicht gefunden werden. Figure 3a**) using blasticidin S deaminase (BSD) as a selection marker. A549 cells were transfected together with a transposase encoding plasmid to achieve a priori stable integration. Following antibiotic selection, we checked for ACE2 expression via Western Blotting and investigated the extent to which A549 cells can be infected in comparison to the highly susceptible Caco2. We infected parental and ACE2 expressing cell lines (named as low and high, based on the expression level) with SARS-CoV-2 isolate FFM1 and 24 h post infection the intracellular RNA and RNA from the cell culture supernatant was isolated and subjected to RT-qPCR (**Supplementary Figure S1**). Subgenomic SARS-CoV-2 RNA (sgRNA) was quantified as a marker of active virus replication, while the virus production of the infected cells was determined by detection of genome copy equivalents using the M gene. Compared to the Caco2 cells, A549 with high ACE2 expression cells could be infected more than four log more efficient with SARS-CoV and two log more efficient with SARS-CoV-2 (**Supplementary Figure S1**). However, in comparison to Caco2, 4 log lower infection efficiency was still observed. For constitutive expression of high levels of TMPRSS2, we used a similar construct with BFP and a hygromycin resistance gene cassette (HygR) with a constitutive TMPRSS2 expression under the control of the EF1alpha promoter (**Figure 3b**). Since both expression constructs carry different fluorescence cassettes (BFP and dTomato) under RPBSA promoter, we sorted cell populations with different expression levels of both receptors using flow cytometry and examined these cell lines for SARS-CoV-2 induced CPE formation. The population with the most distinct CPE was chosen, which, notably, was not the one with the highest expression intensity (Population 2, **Supplementary Figure S2**) and retrieved A549-ACE2^high^-TMPRSS2^low^ cells (A549-AT).

**Figure 3.**
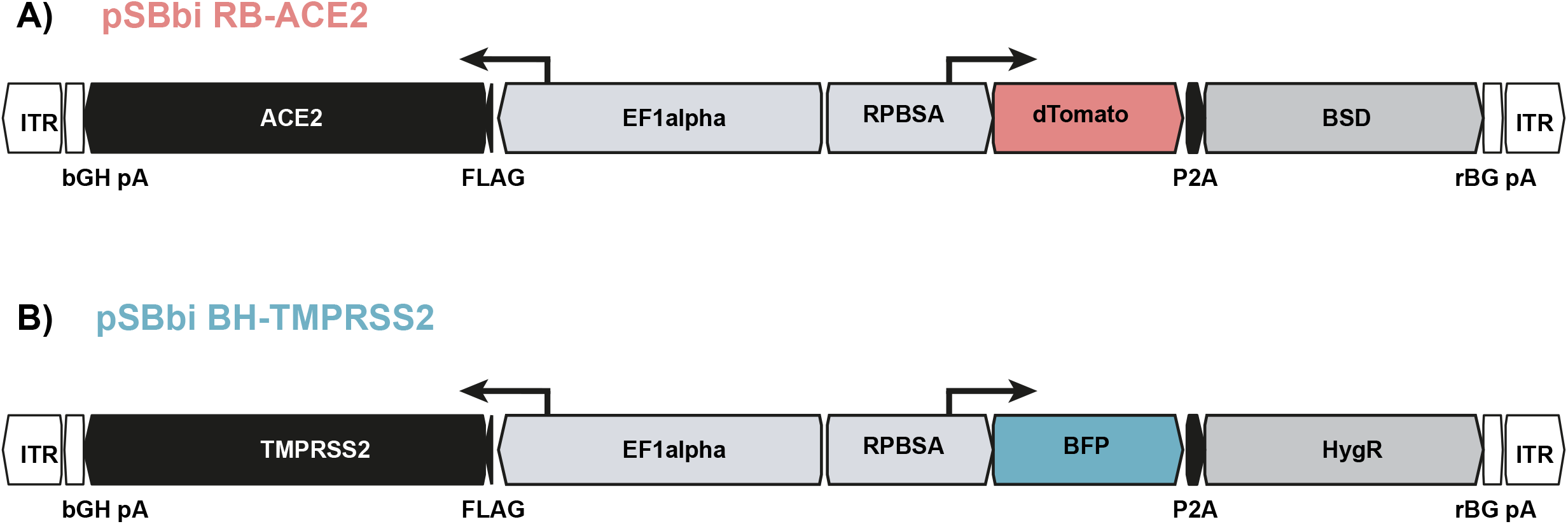
Constructs used for the generation of A549-AT reporter cells. Schematic drawing of plasmids **A)** pSBbi RB-ACE2 and **B)** pSBbi BH-TMPRSS2 including locations of open reading frames (ORFs), transcriptional start sites of the promoters (EF1alpha, RPBSA), inverted terminal repeats (ITR), polyadenylation signals (pA), FLAG epitope (FLAG), 2A self-cleaving peptide (P2A). ACE2 and TMPRSS2 ORFs are marked in black. The ORFs encoding dTomato and BFP are highlighted in red and blue, respectively. The antibiotic resistance cassettes coding for blasticidin S deaminase gene (BSD) and the Hygromycin resistance gene (HygR) are indicated in grey.

**Figure 4.**
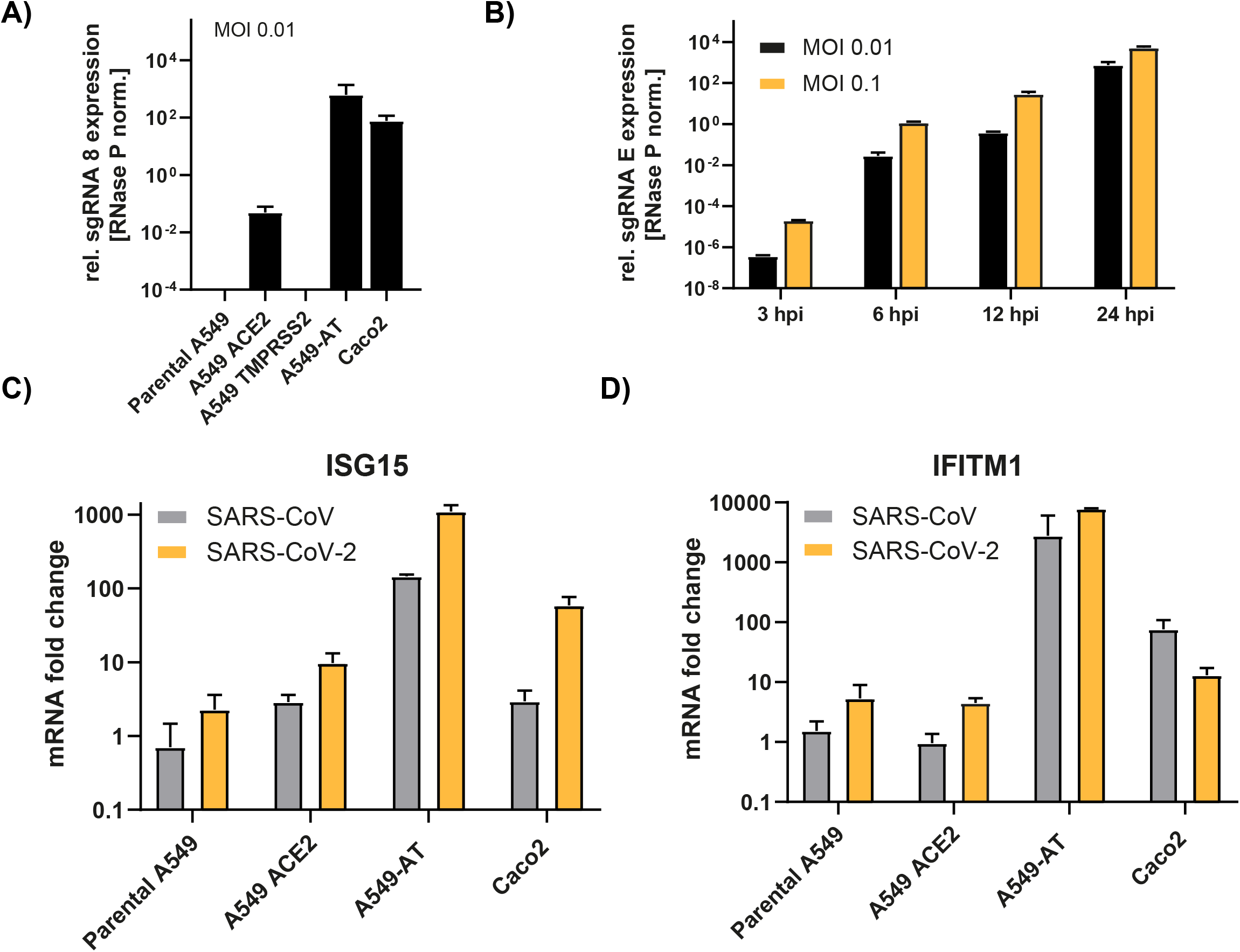
Susceptibility of A549 reporter cells to SARS-CoV-2. Caco2, A549 cells, and derivates were infected with SARS-CoV or SARS-CoV-2 strain FFM1 (MOI as depicted), respectively and total RNA was isolated 24 hours post infection or as indicated. **A and B)** Relative expression of SARS-CoV-2 sgRNA was used as surrogate markers to quantify active viral replication. **C and D)** Relative expression of ISG15 and IFITM1 (fold change) were determined using RT-qPCR to monitor interferon signaling in SARS-CoV and SARS-CoV-2 infected cells.

To compare the properties of the A549-AT reporter cell line with respect to SARS-CoV-2 susceptibility with well-established Caco2 and parental A549 cells, all cell lines were infected with SARS-CoV-2 strain FFM1 (Toptan et al., 2020). 24 h post infection cellular RNA was extracted and subjected to RT-qPCR detecting sgRNA as a surrogate marker for active viral replication. TMPRSS2 overexpression was not sufficient to render A549 cells susceptible to SARS-CoV-2 (**Figure 4A**). Although the expression of ACE2 resulted in a considerable increase in the amount of viral sgRNA, a further increase of 2 log levels was achieved in combination with the additionally introduced TMPRSS2 (**Figure 4A**). This efficiency was thus comparable and even slightly higher when compared to Caco2 cells. Already 3 h after infection, sgRNA species could be detected, with a significant increase measured at 24 h after infection (**Figure 4B**). As expected, the addition of 10-fold more virus (MOI 0.1) resulted in a proportional increase in the sgRNA level. Consistent with **Figure 1**, a marked increase in ISGs (ISG15 and IFITM1) was observed in A549-ATwith both SARS-1 and SARS-CoV-2 (**Figure 4C-D**). The induction of both ISGs was 2 log higher for SARS-CoV and about 1 log higher for SARS-CoV-2 (ISG15) compared to Caco2 cells. IFITM1 expression was induced by 2 log after infection with SARS-CoV and even 3 log for SARS-CoV-2. Taken together, the data show that the newly generated cell line allows robust, time- and concentration-dependent viral replication and induction of strong immune response.

To monitor SARS-CoV-2 replication non-invasively, we used an automated cell imaging system (SparkCyto 400, Tecan). First, we infected the cells with SARS-CoV-2 strain FFM1 (Toptan et al., 2020) and examined the cells for cytopathic effect (CPE) formation. In comparison to the Caco2 cells, we could already observe CPE formation and loss of confluency after 12 hours post infection (**Figure 5A**). In addition, an increasing roughness of the cells could be observed due to the formation of syncytia and cell lysis (Fehler! Verweisquelle konnte nicht gefunden werden. **Figure 5B-D**, **Supplementary Material: Movie 1**). This effect was clearly distinguishable from the non-infected cells (>100 fold) and was proportional to the cell lysis and the associated loss of confluence. Thus, cellular roughness represents an excellent marker for estimating the cytopathic effect, which appears prior to confluence changes. Caco2 cells also showed a clear reduction in confluency and an altered roughness, which however could only be read out automatically to a limited extent in comparison to A549-AT (**Figure 5**, **Supplementary Material: Movies 2 and 3**). The effects were clearly evident at an intermediate MOI of 1, while lower MOIs (0.1 and 0.01) led to a time-delayed onset of roughness and confluency loss. A549-AT were also suitable to display the CPE of various SARS-CoV-2 isolates including variants of concern (VoCs) B.1.1.7 (incl. Spike N501Y) and P.2 (incl. Spike E484K), which have been associated with higher transmission and immune escape, respectively (Widera et al., 2021) (**Figure 5D, G-H**).

**Figure 5.**
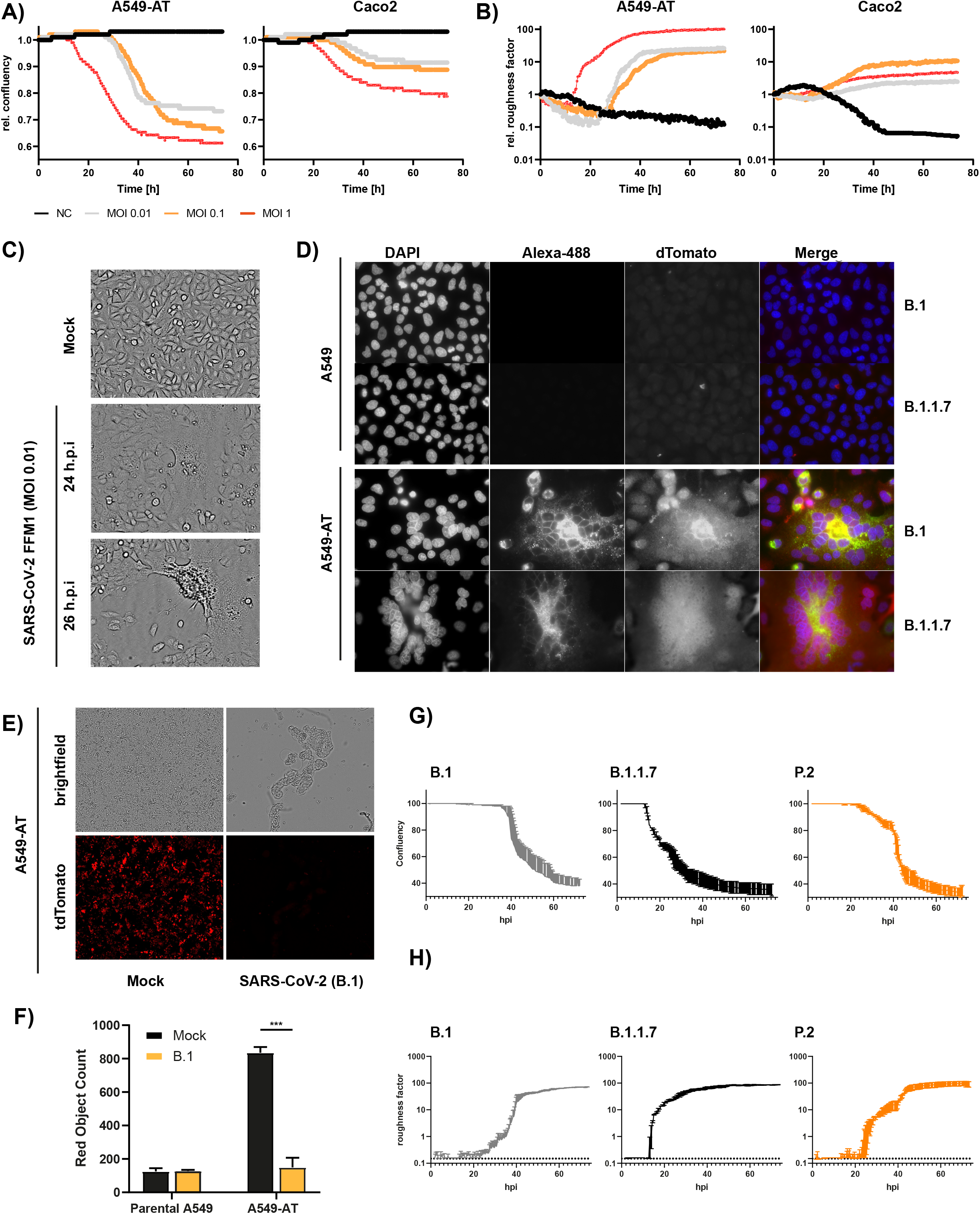
Monitoring of SARS-CoV-2 infection and CPE formation. **A and B)** Caco2 and A549-AT cells were infected with SARS-CoV-2 strain FFM1 with the indicated MOI. Relative confluency and roughness of infected cells was analyzed in an automated live-cell-imaging system in continuous measurement over the specified period in 37°C and 5%CO_2_. **C)** Imaging of SARS-CoV-2 (FFM1) infected A549-AT cells showing induced syncytia and CPE formation. **D)** Fluorescence imaging of SARS-CoV-2 (B.1 and B.1.1.7) infected parental A549 and A549-AT cells. SARS-CoV-2 Spike was stained with an Alexa-488-coupled antibody and cellular dTomato was used to visualize syncytia formation. **E)** Representative fluorescence imaging of A549-AT cells constitutively expressing dTomato enabling the detection of SARS-CoV-2 induced cell lysis. **F)** Red object count was used to quantitate the loss of dTomato fluorescence in SARS-CoV-2 infected cells shown in E). **G-H**). Relative confluency and roughness of cells infected with SARS-CoV-2 variants B.1 (FFM7), B.1.1.7, and P.2. Analysis was performed as described in A).

Quantifying the loss of dTomato signal upon viral infection constitutes a third read-out assay to monitor CPE formation. This is particularly suitable for automated fluorescence intensity measurement, which is a time and resource efficient readout method. The optimal time to monitor fluorescence in SARS-CoV-2 infected cells was at 48 h post-inoculation for low MOI (0.01 and 0.1), and 24 h post-inoculation for high (1 MOI) infectious doses (**Figure 5E-F**). These data demonstrate that the A549-AT reporter cell line is suitable for numerous automated readout methods and for determining SARS-CoV-2 replication. Next, we evaluated to which extent the A549-AT reporter cell line is suitable for drug screenings, testing antiviral drugs and monoclonal neutralizing antibodies (mAbs). For this purpose, we pre-treated cells with increasing amounts of the well described antiviral compound Remdesivir and infected the cells with SARS-CoV-2 strain FFM1 (**Figure 6E**). In agreement with current literature (Wang et al., 2020a), we estimated an IC_50_ of 47.96 nM using relative confluency and roughness.

**Figure 6.**
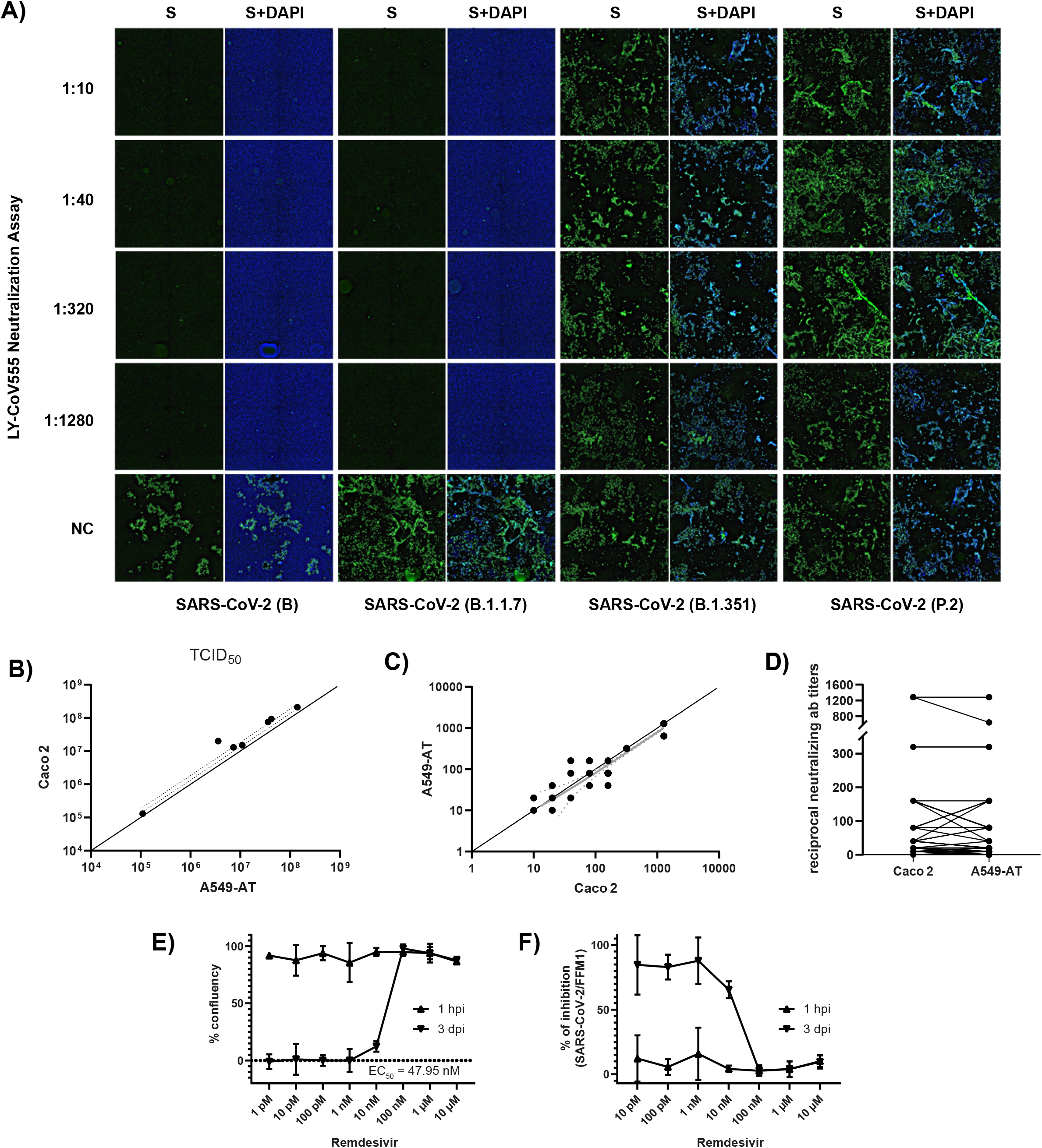
Correlation of neutralization and TCID50 test results obtained with A549-AT and CaCo2 cells. **A)** Neutralization titers against SARS-CoV-2 variants B (FFM1), B.1.1.7, B.1.351, and P.2. The mAb solution containing LY-CoV555 was serially diluted (1:2) and incubated with 100 TCID50 / well. Cells were inoculated and analyzed for a CPE formation after 72 h. Immunofluorescence staining was performed using spike antibody and a secondary antibody coupled to Alexa488 for detection. DAPI was used to stain nuclei. **B)** Correlation of SARS-CoV-2 TCID50 titers determined using A549-AT and Caco2 cells. **C and D)** Reciprocal SARS-CoV-2 microneutralization titers resulting in 50% virus neutralization (NT50). The values indicate mean values from two replicates per cell line. **E and F**) Evaluation of antiviral activity of remdesivir (EC_50_) against SARS-CoV-2. Relative confluency E) and roughness F) were measured as surrogate marker to monitor for SARS-CoV-2 induced CPE formation at the indicated timepoints. Prior to infection, cells were treated with the indicated concentrations of remdesivir.

Next, we determined TCID50 titers of various SARS-CoV-2 isolates and compared the values of A549-AT with Caco2 derived TCID50 titers, which were well correlating (**Figure 6B**). Since mAbs and neutralizing titers of patient derives sera are important factors in SARS-CoV-2 immunity, we used A549-AT cells and determined neutralization ab titers. The reciprocal antibody titers were determined with A549-AT and had an excellent correlation to titers obtained with Caco2 cells (**Figure 6C-D**). SARS-CoV-2 neutralization titer assays showing limiting neutralizing capacity of commercial mAb LY-CoV555 against E484K carrying SARS-CoV-2 isolates (Widera et al., 2021) was also confirmed using immunofluorescence staining. SARS-CoV-2 variant B.1.1.7, as well as the isolates from early 2020 (FFM1 and FFM7), could be efficiently neutralized by bamlanivimab (titer 1/1280, respectively (**Figure 6A**). However, no neutralization effect could be observed against either B.1.135 or P.2, both harboring the E484K substitution. These data highlights that our A549-AT cells are suitable for high-throughput screening for antiviral compounds as well as for investigating the efficacy of neutralizing antibodies against different viral variants.

Since A549-AT was found suitable for automated readout of SARS-CoV-2 infection experiments, viral outgrowth assays as well as drug screenings and neutralization tests, we next focused on the investigation of cellular restrictions and dependency factors. For this purpose, we generated a third vector based on the Sleeping Beauty transposase system, which has a doxycycline (Dox) inducible expression cassette (**Figure 7**). In addition to the gene of interest, which is expressed under the inducible TCE promoter, the constructs also contain a cassette encoding the green fluorescent protein (GFP) and an additional resistance cassette (PuroR) for antibiotic selection. To provide a proof of principle, we cloned the known host restriction factor IFITM1, which has an antiviral effect against SASR-CoV-2, into the expression cassette. A549 and for comparison purposes Caco2 cells were stably transfected with these constructs and subjected to antibiotic selection. As validated with Western blotting, 48 h after doxycycline addition, maximum overexpression of IFITM1 could be achieved (**Figure 7D**). Addition of doxycycline prior to infection resulted in a marked decrease in SARS-CoV-2 sgRNA levels, indicating inhibited virus entry (**Figure 7C**). Of note, the addition of doxycycline itself had no effect on cell viability (**Figure 7E**).

**Figure 7.**
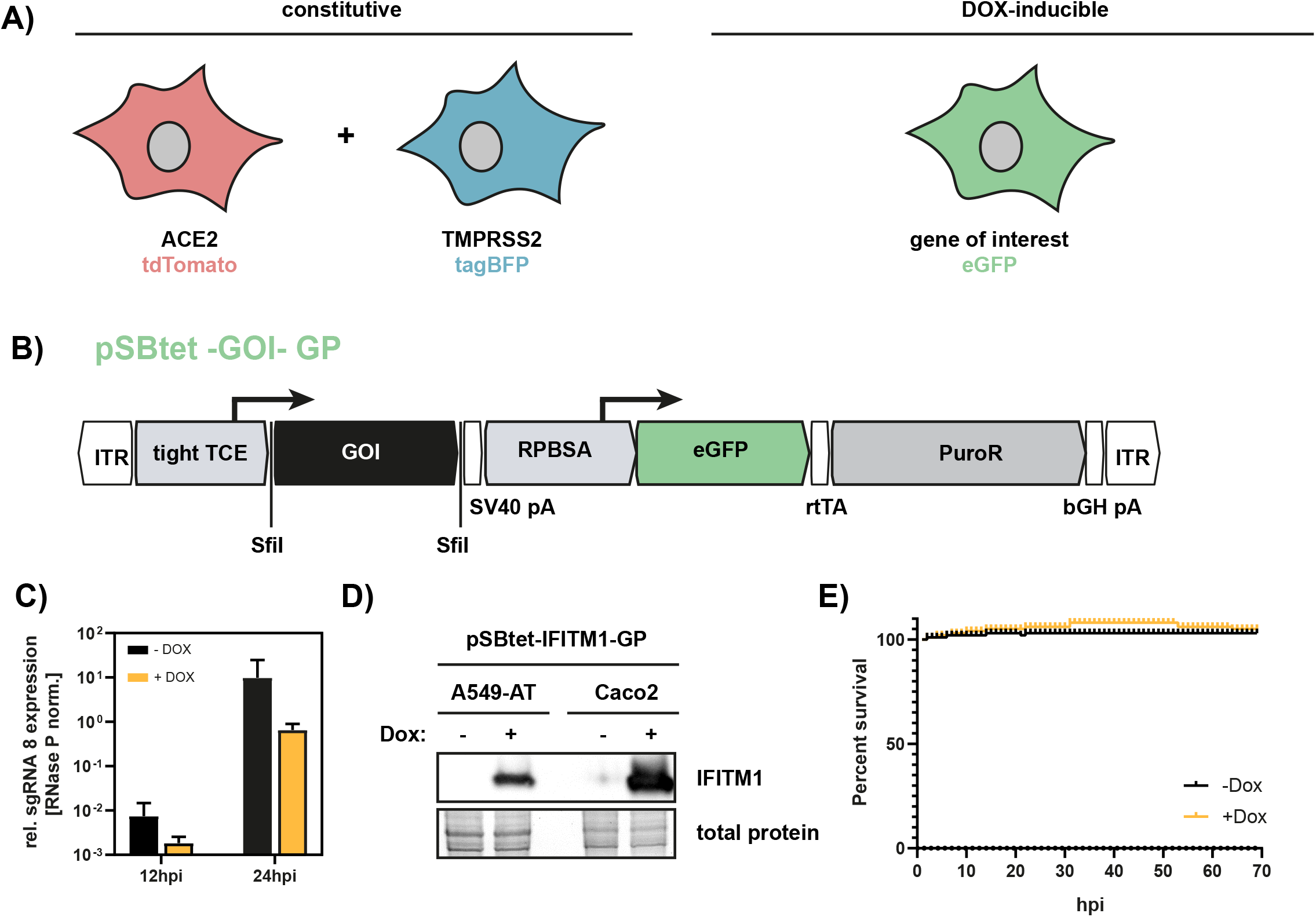
Inducible overexpression of host factors in A549-AT cells. **A)** Graphical overview of A549-AT cells constitutively expressing ACE2/dTomato (red),TMPRSS2/BFP (blue), and doxycycline inducible gene of interest (green). **B)** Schematic drawing of plasmid pSBtet – GOI –GP including locations of open reading frames (ORFs), transcriptional start sites of the indicated promoters (tight TCE, RPBSA), inverted terminal repeats (ITR), polyadenylation signals (pA), SfiI sites for cloning (SfiI), and the reverse tetracycline-controlled transactivator (rtTA). The ORF of the GOI is marked in black. The ORFs encoding eGFP is highlighted in green. The antibiotic resistance cassette encoding the puromycin resistance gene (PuroR) is indicated in grey. **C)** A549-AT cells overexpressing the host restriction factor IFITM1 were infected with SARS-CoV-2 and total cellular RNA was subjected to RT-qPCR analysis at the indicated time points. Relative expression of sgRNA 8 in Dox-induced or mock cells was used as surrogate marker for viral entry and active SARS-CoV-2 replication. **D)** Western blot analysis of A549-AT and Caco2 cells stably transfected with a FLAG-tagged IFITM1 encoding construct illustrated in B. **E)** Cell viability of Dox and mock treated A549-AT cells was determined using a microplate reader with cell imaging and real time – cytometry. Cell viability was analyzed after doxycycline induction measuring cellular confluency.

In summary, by using the Sleeping Beauty transposase system, we generated a robust and versatile reporter cell line that allows SARS-CoV-2 infection experiments compatible for high-throughput and live cell imaging. Sleeping Beauty transposase-based overexpression and shRNA based knockdown cell lines with a second constitutive fluorescence reporter cassette (eGFP) can be generated in a very short time, enabling the investigation of host and restriction factors in a doxycycline -inducible manner.

## 4. Discussion

SARS-CoV-2 research studies require appropriate cell culture model systems. However, the current methods to monitor and quantitate SARS-CoV-2 infectivity in cell culture are time-consuming and labor-intensive. The A549-AT system established in this work allows to study the interactions between SARS-CoV-2 and the host cell in a high throughput setup at low cost. In addition, assays common in antiviral research can be transferred to an A549-AT setting with comparable results, but in less time.

The cell line A549 was established in 1972 by D.J. Giard, et al. by culturing an explant of lung carcinoma tissue from a 58-year-old male Caucasian patient (Giard et al., 1973). The alveolar adenocarcinoma cell line (alveolar type II-like) combines a lung-associated background with the advantages of an easy-to-handle cell system. In our study, we compared the properties of the A549 cell line to Caco2 well established in the field and in our lab (Bojkova et al., 2020b; Shin et al., 2020; Toptan et al., 2020) and found clear advantages in terms of proliferation, trypsinization and transfection efficiency (**Figure 2**). In addition, A549 cells can be subjected to efficient antibiotic selection, as significantly lower concentrations and treatment times are required when compared to Caco2 cells (data not shown). Another advantage become apparent during microscopic observation, as Caco2 cells are a very heterogeneous cell line (Lea, 2015), while A549 have a much more homogeneous morphology (**Figure 2**).

Cellular entry of coronaviruses is a multistage process involving virus attachment to the cell surface, receptor engagement, protease processing and membrane fusion (Li et al., 2021). These stages rely on distinct domains in the SARS-CoV-2 spike protein (Lan et al., 2020). Emerging SARS-CoV-2 variants of concern (VOCs) like B.1.1.7, B.1.351, P.1, P.2, B.1.672 initially observed in the United Kingdom, South Africa, Brazil, and India, respectively, harbor amino acid substitutions in the receptor binding domain of the Spike protein (Garcia-Beltran et al., 2021; Hoffmann et al., 2021). First and foremost, N501Y was described to increased infectivity due to enhanced receptor binding (Starr et al., 2020). Further studies have shown that E484 represents an immunodominant site on the RBD since E484K reduced naturalization capacity of human convalescent sera by >100-fold (Greaney et al., 2021; Widera et al., 2021). These amino acid substitutions have also been observed to evade the antibody response elicited by an infection with other SARS-CoV-2 variants, or vaccination. In this work we could show that A549-AT were susceptible to multiple variants allowing studies with a broad range of viral strains. In addition, mutagenesis of ACE2 or TMPRSS2 and subsequent phenotypical characterization can be performed with our system

Through the stable integration of fluorescent proteins, in particular the constitutive dTomato expressing cassette localized on the ACE2 expression vector (**Figure 3**), the reporter cell line enables fast and automatic readout using red fluorescence for CPE and confluency determination. By using the additional Dox-inducible system (pSBTet-GP vector backbone), also cellular host factor can be overexpressed in order to study and dissect SARS-CoV-2 biology. All successful transfection experiments can be easily monitored by the expressed color tags which allow also the automation of all subsequent screening experiments.

In addition to the loss in fluorescence, two non-invasive factors can also be used as surrogate markers for CPE formation and cell lysis, which are confluency and roughness. The former was proofed as highly robust and was additionally confirmed with DAPI staining (data not shown). Exclusively in A549-AT, cellular roughness was used to monitor syncytia formation which was much less pronounced in Caco2 cells. Furthermore, a clear CPE formation could be monitored since A549 have a less heterogenic morphology than Caco2 (**Figure 5**).

An important factor to study virus-host cell interactions is the ability to physiological respond to external stimuli and to induce a physiological antiviral response. Thus, we tested the interferon signaling pathways by treatment with interferon types I, II, and III (IFNs I-III). The IFN-I signaling deficiency in Vero cells allows several viruses to replicate efficiently. However, to study physiological responses to viral infections, intact interferon signaling is mandatory. In our study and in line with other studies (Li et al., 2021), we observed the induction of the type I interferon-induced genes ISG15 and IFITM1 correlating with the efficiency of virus entry (**Figure 3**). Consistently, insertion of an ACE2 expression cassette to parental A549 cells led to a small increase in ISG15 expression after SARS-CoV-2 infection, which was significantly increased by additional insertion of TMPRSS2. This increase correlated with the amount of detected sgRNA, which is a surrogate marker of active viral replication (Kim et al., 2020; Wolfel et al., 2020), and was higher in A549 cells when compared to SARS-CoV-2 infected Caco2 cells. Noteworthy, in both cell lines we observed higher ISG15 levels after SARS-CoV-2 infection compared to SARS-CoV infection, which was in line with previously published data showing a higher interferon antagonizing capacity of SARS-CoV (Shin et al., 2020).

In humans, type I interferons consist of IFN-α, β, (ε, κ, and ω), while especially IFN-α consists of multiple subtypes with distinct biological activities (Lavender et al., 2016). While type I interferons are mostly produced by infected cells and cells of the immune system, type II interferon (IFN-γ) is predominantly produced by immune cells. In our study IRF1 expression was used as a surrogate marker of IFN-II induction and was induced in Vero, Caco2 and A549 cells to a comparable extent. Importantly, IFN-γ driven inflammation was shown to induce a vulnerable state in vivo allowing SARS-CoV-2 replication by promoting ACE2-expression and enhanced virus production in infected cells (Heuberger et al., 2021). Hence, A549-AT cells are suitable to study cell culture based responses to IFN-γ.

Type III interferons consist of four subtypes of IFN-λ. Although some immune cells produce type III interferons, these are also produced by epithelial cells or cells that are from the same developmental stage, which are the respiratory tract, the gastrointestinal tract, and the urogenital tract derived cells. Type III IFNs were effective in preventing SARS-CoV-2 infection but are in contrast to IFN-I locally limited in the respiratory tract (Andreakos and Tsiodras, 2020), which makes them relevant for treatment in early infected patients. Hence, it will be also interesting to analyse the impact of type III interferons on different SARS-CoV-2 variants in future.

Although lentiviral transduction is relatively quick, the compatibility of a gene of interest, which has to be cloned in a sophisticated vector, is size restricted and does not allow polyA and splicing sites, which might interfere with the LTR-mediated integration of the transgene into the host chromosome. Random integration into genome preferentially in transcriptionally active regions might cause cell cycle defects. Depending on several factors a varying copy number of integrations might occur. Furthermore, the preparation of lentiviruses requires S2 Lab work. In contrast, transposable elements as the Sleeping Beauty (Kowarz et al., 2015) offers several advantages for transgene integration. The use of SfiI with distinct recognition sequences at the 5’ and 3’site allows directional cloning with only one enzyme, which is rarely found in GOIs due to the long recognition sequence. In comparison to other methods Sleeping Beauty Transposase is quick and might be performed under S1 safety labs. Outstanding features are no size restriction, compatibility with the integration of polyA and splicing sites in the gene of interest region. Integration into genome occurs preferentially at TA-sites, however a certain limitation remains since a targeted integration at a predetermined location in the genome is not possible. Through precise dosing of the vector and the transposase-encoding plasmid scalable number of integrations can be achieved (Skipper et al., 2013).

A further conceivable development of the system would be a construct expressing eGFP enabling the parallel expression of an inducible shRNA-mediated knock-down cassette. This would allow loss-of-function experiments to be carried out in addition to gain-of-function experiments.

In conclusion, we developed a reporter cell system with optimal ACE2/TMPRSS2 ratio for high SARS-CoV-2 susceptibility, syncytia and CPE formation. The confluency and particularly roughness of A549-AT cells represents a highly sensitive marker for early syncytia formation and cytopathic effects. Thus, A549-AT cells allow the infection and readout of an experiment on the same working day. Using 1 MOI the system theoretically allows infection and read-out within one working day performed via non-invasive roughness factor analysis.

## Acknowledgements

We thank Marco Bechtel for helping to establish ACE2 Western Blotting, S. Stein and A. Trzmiel for cell sorting of stably transfected fluorescent A549 cells, Wibke Ballhorn for her support during fluorescence microscopy. We are thankful for the numerous donations to the Goethe-Corona-Fond and for the support of our SARS-CoV-2 research. M.W. and R.M were supported by the Deutsche Forschungsgemeinschaft (DFG, WI 5086/1–1, DFG MA 1876/13-1). All authors have read and agreed to the published version of the manuscript.

## Materials and methods

### Cell culture and virus propagation

Vero, A549 and Caco2 cells were cultured in Minimum Essential Medium (MEM) supplemented with 10% fetal calf serum (FCS), 2% L-glutamine, 100 IU/ml of penicillin and 100 μg/ml of streptomycin at 37°C and 5% CO_2_.

Cell free virus aliquots of SARS-CoV isolate FFM1 (Drosten et al., 2003) and SARS-CoV-2 isolates FFM1, FFM7, B.1.1.7, B1.351, P.2 (Hoehl et al., 2020; Toptan et al., 2020; Widera et al., 2021) were stored at −80°C and titers were determined by TCID50 in Caco2 cells. According to the Committee on Biological Agents (ABAS) and Central Committee for Biological Safety (ZKBS) all infectious work was performed under biosafety level 3 (BSL-3) conditions. Efficiency of virus inactivation were confirmed as previously published (Westhaus et al., 2020).

### Interferon stimulation

Vero, Caco2 and A549 cells were stimulated with the type I-III interferons (500 U / mL and 10 ng / mL, respectively. Total RNA was isolated and subjected to RT-qPCR analysis at the indicated time points.

### Plasmids construction and cloning

Total RNA (1 μg) was isolated from HEK293T cells and reverse transcribed into cDNA using random hexamers and SuperScript II RT (Invitrogen). The ACE2 and TMPRSS2 genes were amplified with GoTaq polymerase (Promega) by using the oligonucleotides ACE2.Flag.Sfi.F, ACE2.Sfi.R, TMPRSS2.Flag.Sfi.F and TMPRSS2.Sfi.R, respectively (**Supplementary Table 2**). All 3’-primers encode an additional N-terminal FLAG-tag as well as the 5’-SfiI-restriction site, while the 5’-primers encode the additional 3’-SfiI-restriction site. All amplimers were cloned into the PCR2.1-Topo-TA vector (Invitrogen) for Sanger sequencing. Correct amplimers were digested with Sfi1 and cloned into the pSBbi-RB (ACE2) or pSBbi-BH (TMPRSS2) Sleeping Beauty plasmids (Kowarz et al., 2015).

### Generation of stable cell line

Using TransIT LT1 (Mirus) A549 and Caco2 were transfected with a transposase-encoding plasmid (SB100x) and the respective Sleeping Beauty construct (see above). 48 h later, appropriate selection was carried out with hygromycin and blasticidin. A total of three rounds of selection were performed, each with a recovery phase lasting two doublings per cell line. Transfection efficiency was monitored by fluorescence microscopy using the fluorescent proteins dTomato and mTag-BFP.

### Quantitative proliferation and trypsinization assay

CaCo2 and A549-based cells were seeded in 12-well plates (2×10^4^ A549-AT and 1×10^5^ Caco2 / well, respectively). Cells were incubated for 16 days at 37°C and 5% CO_2_ and cell count and confluency was optically determined using SparkCyto 400 (Tecan) at regular intervals starting after 48 hours after seeding. To allow attachment, cells were incubated for 48 h before monitoring. Prior to imaging the wells were washed using PBS and refilled with 2 ml culture medium. Trypsinization assay was performed in 96 well plates. After rinsing twice with PBS, confluent cells were treated with 100 μl trypsin (2 mg/ml) and incubated for different intervals. Cells were incubated at room temperature and 37°C for incubation times longer than 10 min. Subsequently cells were rinsed with cell culture medium, remaining cells were fixed with 3% PFA, and nuclei were stained with DAPI and analyzed with a plate reader (SparkCyto 400).

### RT-PCR and RT-qPCR analysis

SARS-CoV-2 RNA from cell culture supernatant samples was isolated using AVL buffer and the QIAamp Viral RNA Kit (Qiagen) according to the manufacturer’s instructions. Intracellular RNAs were isolated using the RNeasy Mini Kit (Qiagen) as described by the manufacturer. For detection of intracellular and extracellular SARS-CoV-2 genomic RNA, primers and dual-labeled probes were used **(Supplementary Table 1**). Multiplex RT-qPCR assay were carried out using Reliance One-Step Multiplex RT-qPCR Supermix (BioRad) or LightCycler Multiplex RNA Virus Master (Roche). SYBR green based RT-qPCRs were performed using Luna Universal One-Step RT-qPCR Kit (NEB).

### Western blot analysis

Protein extracts, separated by SDS-PAGE and transferred onto PVDF membranes, were probed with antibodies against FLAG (F1804, Sigma), ACE2 (ab15248, Abcam) or GAPDH (2275-PC-100, Trevigen). Proteins of interest were detected with HRP-conjugated secondary IgG antibody and visualized with ECL Western blotting substrate (Thermo Scientific) according to the provided protocol.

### SARS-CoV-2 Neutralization Assay

The mAb solutions were serially diluted 1:2 and incubated with 4000 TCID50 / ml of each SARS-CoV-2 variant and subjected to cell based SARS-CoV-2 neutralization assay (Widera et al., 2021). The corresponding sample dilution resulting in 50% virus neutralization titer (NT50) was determined. After 3 days of incubation cells were evaluated for the presence of a cytopathic effect (CPE). Spike Staining was performed as described below.

### Microscopy and imaging of SARS-CoV-2 infected cells

Time-lapse images were taken with the SparkCyto 400 multimode reader with integrated microscope unit (TECAN). The cell culture condition was maintained with the integrated incubation chamber with 5% CO_2_ and 37°C. Cells were imaged in regular intervals as indicated. Confluency and roughness measurements were performed with the 10x objective. A single central image was used to record movies showing the cytopathic effect (**supplementary data, Movies 1 - 3**). Red object count, Confluency and roughness were analyzed using ImageAnalyzer Software v.1.1 (Tecan). Immunofluorescence experiments were performed using SARS-CoV and SARS-CoV-2 Spike antibody (40150-R007, Sino Biologicals) and Alexa-488 conjugated anti-rabbit secondary antibody (Invitrogen).

### Statistical analysis

If not indicated differently all infection experiments were repeated in three independent experiments. Statistical significance compared to untreated control was determined using unpaired student’s t-test. Asterisks indicated p-values as * (p < 0.05), ** (p ≤ 0.01) and *** (p ≤ 0.005).

