## Supplementary figures and images for "Generation of a Sleeping Beauty transposon-based cellular system for rapid and sensitive identification of SARS-CoV-2 host dependency and restriction factors"

### Supplemental Figure 1

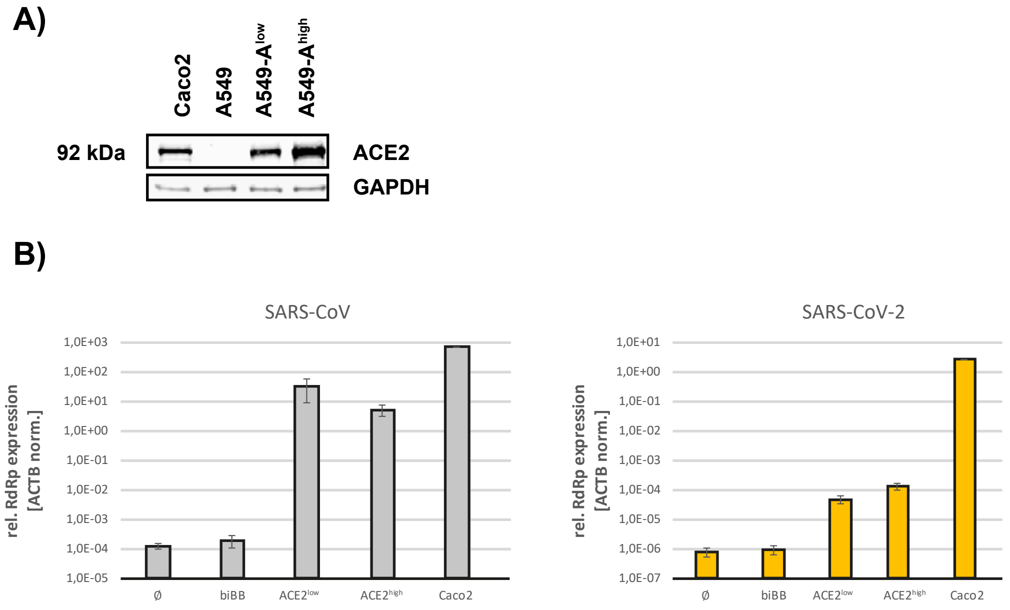
