## Supplemental Figure 2 for "Generation of a Sleeping Beauty transposon-based cellular system for rapid and sensitive identification of SARS-CoV-2 host dependency and restriction factors"

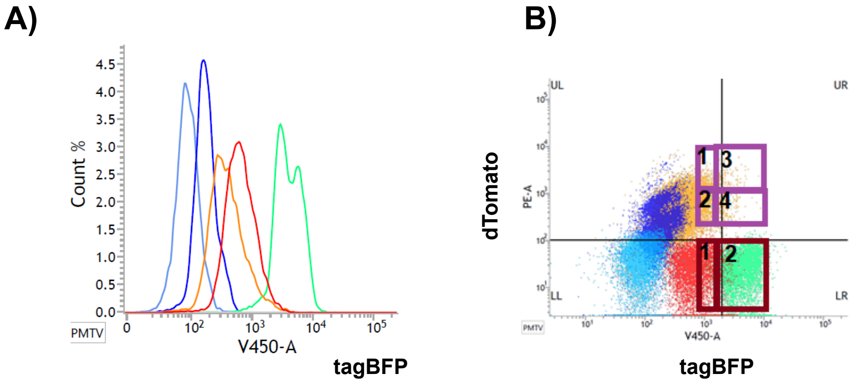


**Population 1: A549-ACE2$^{veryhigh}$ + TMPRSS2$^{low}$**
**Population 2: A549-ACE2$^{high}$ + TMPRSS2$^{high}$**
**Population 3: A549-ACE2$^{veryhigh}$ + TMPRSS2$^{high}$**
**Population 4: A549-ACE2$^{high}$ + TMPRSS2$^{high}$**
